# Dysregulation of LINGO1 expression in the SOD1(G93A) mutant mice during symptomatic stages of the disease and early postnatal oligodendrogenesis

**DOI:** 10.64898/2026.09.03.749143

**Authors:** Amina Zahaf, Julie Bourseguin, Laetitia Cobret, Abdelmoumen Kassoussi, Mireille Moussaed, Cédric Raoul, Elisabeth Traiffort, Séverine Morisset-Lopez

**Author notes:** Corresponding authors: Séverine Morisset-Lopez: Address: Centre de Biophysique Moléculaire, UPR 4301 CNRS, Orléans University, rue Charles Sadron, CEDEX 02, 45071 Orleans, France, Elisabeth Traiffort: INSERM-Paris Saclay University, Diseases and Hormones of the Nervous System U1195, 80 Rue du Général Leclerc 94276 Le Kremlin-Bicêtre, France. Author contributed equally to this work (Amina Zahaf, Julie Bourseguin, Laetitia Cobret).

## Abstract

Oligodendrocytes play an essential role in axonal metabolic support and myelination, and their dysfunction is associated with amyotrophic lateral sclerosis (ALS). LINGO1 is a major inhibitor of oligodendrocyte differentiation and axonal regeneration, identified as a promising therapeutic targeted for inflammatory demyelinating diseases. Here, we investigated the expression pattern of LINGO1 in the well-established SOD1(G93A) transgenic mouse model of ALS. We show that LINGO1 protein expression is significantly elevated in both grey and white matter of the mutant spinal cord during both the symptomatic and end stages of the disease. LINGO1 is also notably co-expressed with astroglial and neuronal markers. However, neither the levels of LINGO1 transcripts, nor levels of microRNAs predicted to regulate its expression were altered in the mutant spinal cord, suggesting the involvement of alternative regulatory mechanisms. Our analysis of microRNAs revealed in the mutant a significant downregulation of miR-138, a microRNA known to promote myelination. The differential expression of LINGO1 and miR-138 in ALS mice thus may drive deleterious processes during disease progression in adulthood. Moreover, our data show that white-matter pathology arises earlier than initially thought, as evidenced by significant alterations of oligodendrogenesis that manifest as early as postnatal day 5. Study of the molecular determinants of this developmental defect shows increased miR-138 expression, which may promote oligodendrocyte progenitor cell maturation and involve compensatory mechanisms. Overall, our work identifies disrupted expression of LINGO1 and/or miR-138 in brain during early oligodendrogenesis and at symptomatic disease stages. These findings open the way to further investigations that may consider miR-138 as a potential presymptomatic biomarker and LINGO1 as a therapeutic target to promote remyelination in the context of ALS.

## Introduction

Amyotrophic lateral sclerosis (ALS) is a progressive neurodegenerative disease characterized by the selective loss of motor neurons in the brain and spinal cord, leading irreversibly to paralysis and death within 3 to 5 years after symptom onset. Besides riluzole, which is the most commonly used anti-glutamatergic medication, few treatments are in clinical trials, primarily focusing on skeletal muscle, energy metabolism, or cell replacement [1, 2]. More recently, the antisense oligonucleotide tofersen that targets SOD1 mRNA, has been approved for SOD1-associated ALS, offering a precision approach for genetically defined patient [3]. However, although these therapies offer some benefits, there is still no cure or universally effective treatment available.

Despite upper and lower motor neurons are the primary target in ALS, the disease has also been associated with oligodendrocyte (OL) abnormalities. Pathological inclusions have been observed in OL in post-mortem tissues and murine models [4–7]. OL degeneration often precedes motor neuron loss, as shown in the SOD1(G93A) mice, in which deletion of the mutant SOD1 from OLs significantly delays disease onset [4]. In accordance with this finding, neurodegeneration secondarily triggered by OL pathology can also be observed in most demyelinating diseases of the central nervous system (CNS), such as multiple sclerosis [8]. However, while OL dysfunction and demyelination may together drive axonal degeneration, the latter also likely exacerbate OL dysfunction [9].

Remarkably, in sporadic and familial ALS, gray matter demyelination was mainly reported in CNS regions where motor neurons locate [10, 11]. The CNS has the capacity to spontaneously repair the disrupted myelin sheaths via the recruitment, proliferation and differentiation of oligodendrocyte progenitor cells (OPCs), scattered throughout the CNS. In ALS, OPCs are efficiently mobilized and differentiate into OLs. However, these newly formed OLs fail to remyelinate axons, thereby exacerbating motor neuron injury [4, 6]. Thus, therapeutic approaches aimed at promoting OL survival or enhancing their metabolic support to neurons could be helpful for preventing motor neuron degeneration in ALS and potentially mitigating OL dysfunction [4, 6, 12, 13].

LINGO1 is a CNS-specific transmembrane protein expressed in OLs and neurons that regulates neuronal survival, axonal regeneration, and OPC differentiation [14, 15]. LINGO1 mediates inhibitory signaling by interacting with myelin-associated proteins such as Nogo-A, MAG, and OMgp and by forming complexes with NgR1, p75 or TROY [16]. Notably, Nogo-A was shown to be upregulated in ALS muscle and to destabilize neuromuscular junctions [17–19]. Anti-Nogo-A antibodies nevertheless failed in ALS trials [20], suggesting that Nogo-A blockade alone is insufficient. However, targeting LINGO1 and its signaling partners remains a promising strategy. Indeed, experimental models have demonstrated that LINGO1 expression by OPCs and/or axons inhibits oligodendroglial differentiation and myelination. Conversely, LINGO1 inhibition promotes myelination, axonal growth, and neuronal survival [15, 21]. These findings positioned LINGO1 as a therapeutic target for demyelinating diseases, such as multiple sclerosis. The clinical development of the humanized anti-LINGO1 antibody opicinumab provided the first translational evidence of remyelination in humans. In the RENEW trial for acute optic neuritis, opicinumab treatment improved visual evoked potential latency compared to placebo, suggesting functional myelin repair [22, 23]. However, subsequent multiple sclerosis trials, such as SYNERGY, failed to meet their primary endpoints, highlighting the challenge of promoting remyelination across diverse lesion types [24].

MicroRNAs (miRNAs) are small non-coding RNAs that post-transcriptionally regulate gene networks implicated in neurodegeneration. Growing evidence links their dysregulation to ALS [25]. miRNAs also regulate OL development and may serve as biomarkers or therapeutic targets [26]. For instance, in ALS mice, miR-206 upregulation is associated with a delay in disease progression and promotion of neuromuscular synapse regeneration [27]. The expression of miRNAs miR-206, 143-3p and 374b-5p is dysregulated in the serum of ALS patients [28]. Recent meta-analyses confirm consistent dysregulation of certain miRNAs across studies, though variability in cohorts and methodologies limit their direct clinical use [29]. Therapeutically, the broad miRNA field has reached human trials in non-ALS indications, providing evidence for the possible use of miRNA mimics/antagonists while leading to invaluable information for future ALS-directed interventions [30].

According to the regulatory role of miRNAs in OL development and maintenance of myelin integrity [31], and the central role of OL dysfunction in ALS, targeting LINGO1 and miRNA that regulate its expression or myelination could offer new therapeutic avenues. The present study aimed to determine whether LINGO1 dysregulation can be observed in an ALS animal model (SOD1 mice) and whether miRNAs controlling LINGO activity or myelination are altered, with the ultimate goal of identifying innovative therapeutic strategies for ALS.

## Materials and methods

### Animals

SOD1 (G93A) mice were purchased from the Jackson Laboratory (B6.Cg-Tg(SOD1*G93A)1Gur/J ; #004435). All animals were housed in standard conditions: ambient temperature at 20°C, relative humidity at 45–65%, 12h light-dark cycle with food and water ad libitum. All procedures were performed according to the European Communities Council Directive (86/806/EEC) for the care and use of laboratory animals and were approved by the Regional Ethics Committee CEEA26, Ministère de l’Education Nationale, de l’Enseignement et de la Recherche. For immunostaining experiments, adult female mice were transcardially perfused with 4% paraformaldehyde (PFA) in phosphate-buffered saline (PBS). Spinal cords with vertebrae were dissected from pre-symptomatic (postnatal day 75), symptomatic (P110), and end-stage (P135) SOD1 (G93A) mice, as well as age-matched wild-type (WT) controls. Samples were post-fixed in 4% PFA for 24 hours at 4 °C, dehydrated through graded alcohol baths, embedded in paraffin, and sectioned at 7 μm using a rotary microtome. Sections were subsequently processed for immunostaining.

Both male and female pups at postnatal days 0 (P0) and 5 (P5) were deeply anesthetized, and the brains were rapidly removed and fixed in 4% PFA overnight at 4 °C, cryoprotected sequentially in 10%, 20%, and 30% sucrose in PBS. After cryoprotection, tissues were embedded, frozen, and sectioned at 14 µm using a cryostat for immunostaining. For RT-PCR experiments, spinal cord derived from both male and female mice at pre-symptomatic (postnatal day 75), symptomatic (P110), and end-stage (P140) SOD1 (G93A) as well as age-matched wild-type (WT) controls were dissected and snap-frozen in liquid nitrogen. Samples were then stored at −80°C until further processing. Brains derived from both male and female SOD1 (G93A) and WT mouse pups at P0, P5 and P8 were similarly processed.

All animal research has been performed according to the ARRIVE guidelines [32]. Due to the small litter sizes and the short time window for animal reproduction, the data obtained did not allow us to address the question of possible sex-related differences at all symptomatic stages. However, this question will be likely important to address in the context of future studies.

### Protein extraction

Proteins were extracted in a lysis buffer (50mM Tris pH 7.5, 150mM NaCl, 10mM EDTA, 1% Triton X100) containing protease inhibitor cocktail (Roche, 1187358001). Pulse sonication was performed on ice and applied 4×5sec ON, 4×5sec OFF. Then, the lysates were centrifuged at 10 000x g for 10 min at 4°C. Relative protein amounts were quantified with DC Protein Assay Biorad. Laemmli buffer 4X (200mM Tris-HCl pH6.8, 4% SDS, 40% glycerol, 0.02% bromophenol and 0.5M β-mercaptoethanol) was added to the supernatant.

### Western blotting analysis

Proteins were separated on 10% Tris-glycine SDS/PAGE and transferred onto PVDF membranes (GE Healthcare Life Sciences). The membranes were blocked 1hour at RT with 5 % non-fat dry milk in PBS1X 0.1% Tween20. Membranes were incubated in the same buffer with primary antibodies overnight at 4°C with agitation. The following antibodies were used: anti-Lingo1 (abcam, ab23631, 1:1000 / R&D, AF3086 1:500), anti-Actin (cell signalling, #4970S, 1:1000), anti-GAPDH (cell signaling, #5174, 1:1000). Horseradish-peroxidase-conjugated Goat anti-Rabbit IgG (H+L) (Invitrogen, # 65-6120, 1:40 000) and Rabbit anti-Goat IgG (H+L) (Invitrogen, # 81-1620, 1:40 000) were used as secondary antibodies, 1hour at RT. Immunoreactive bands were detected using the ECL detection kit (Biorad, 170-5060), with an Imaging System (Biorad, Chemidoc).

### Immunostaining experiments

The primary antibodies were as follows : mouse anti-Adenomatous Polyposis Coli (APC/CC1; OP80; 1:500; Calbiochem), mouse anti-Glial Fibrillary Acidic Protein (GFAP; G3893; 1:1500; Sigma-Aldrich), rat anti-Platelet-Derived Growth Factor Receptor alpha (PDGFRα; 558774; 1:500; BD Pharmingen), mouse anti-Ki67 (550609; 1:150; BD Pharmingen), rabbit anti-LINGO1 (Ab2363; 1:250; Abcam), and goat anti-Choline Acetyltransferase (ChAT; Ab144P; 1:500; Sigma-Aldrich). Secondary antibodies were : goat anti-mouse Alexa Fluor 488 (A11029; 1:250; Thermo Fisher Scientific), goat anti-rat Alexa Fluor 633 (A21094; 1:750; Thermo Fisher Scientific), goat anti-mouse Alexa Fluor 546 (A11030; 1:500; Thermo Fisher Scientific), goat anti-rabbit Cy3-conjugated (111165003; 1:250; Jackson ImmunoResearch), donkey anti-goat Alexa Fluor 488 (A11055; 1:500; Thermo Fisher Scientific), and donkey anti-rabbit Alexa Fluor 555 (A31572; 1:750; Thermo Fisher Scientific).

### Imaging and analysis

Images were acquired with Axiovision 4.2 (Carl Zeiss) epifluorescence microscope and analyses performed with Fiji software. A total of 3–5 sections per mouse were analyzed. The immunofluorescent-positive cells or areas were determined in one every other 5 sections throughout the developing corpus callosum or the adult lumbar spinal cord per mouse and then averaged for each animal. Cell density was manually quantified on x20 images using the CellCounter plugin. Fluorescent area quantification and colocalization analyses were performed using ImageJ. For area measurements, all images were converted to 8-bit and processed identically across experimental conditions. A fixed threshold was applied uniformly, and the area of regions of interest was quantified using standard measurement tools. Colocalization was assessed using the Image Calculator function with the “AND” operation applied to the two channels, generating an image containing only overlapping pixels. This image was thresholded using identical parameters across all conditions, and the area displaying colocalized fluorescent signals was quantified.

### RNA extraction and Quantitative RT-PCR

RNA was extracted using an miRNeasy Tissue/Cells Advanced minikit as per the manufacturer’s instruction (Qiagen 217604). RNA was treated with DNAse I (Thermofisher). Disruption and homogenization were performed using the Tissue Ruptor gentle MACS octo Dissociation (Miltenyi Biotec). RNA integrity was verified with Agilent RNA 6000 NanoChips, according to the manufacturer’s instructions, then reverse transcribed using Maxima First Strand cDNA Synthesis kit (Thermo Scientific, K1641/1642) or miRCury LNA RT Kit (Qiagen 339340) according to the manufacturer’s instructions. For oligonucleotide sequences, we used commercially available primers for Lingo1 (RT^2^ Qiagen, PPM34752A and quantitect Qiagen, QT00147322), Ppia (quantitect Qiagen, QT00247709), Hprt1 (quantitect Qiagen, QT00166768), Gapdh (quantitect Qiagen, QT01658692), miR138-2-3p (RT^2^ Qiagen, YP02111611), miR-23b-5p RT^2^ Qiagen, YP02108273, miR33-5p RT^2^ Qiagen, YP00205690, miR140-3p1 RT^2^ Qiagen, YP00204304; miR184-5p (RT^2^ Qiagen YP02113251); miR219-5p RT^2^ Qiagen, YP02117691).

RT-qPCR was performed using ONEGreen FAST qPCR Premix kit (Ozyme OZYA 008-40) or miRCURY LNA SYBR Green PCR (Qiagen, 339345) with miRCURY LNA miRNA PCR assay (Qiagen, 339306) according to the manufacturer’s instructions and the LightCycler 480 instrument (Roche). The data were analysed using LightCycler softward (Roche). The following cycling program was used : initial denaturation and polymerase activation at 95°C for 2 min; 40 cycles with 5s at 95°C followed by 30s at 60°C ; followed by a melting curve step at 15s at 95°C. Then, 1min at 60°C, 15s at 95°C and 4°C unlimited. Cycle threshold (Ct) values for all mRNAs of interest were determined for three technical replicates per sample, normalized to Ppia, Hprt1, Gapdh reference mRNA, and the data were expressed as fold-change relative to the control using the 2^-ΔΔCt^ method.

### Statistical analysis

Statistical analysis was performed with GraphPad Prism 7.0 software (La Jolla, CA). The significance of differences between means was evaluated by ANOVA followed by Tukey’s or Holm-Sidak’s post-tests for comparisons. The values are the means ± SEM from the number of animals indicated in each plotted graph or as indicated in the corresponding legends.

## Results

### LINGO1 is increased at different stages of the disease in SOD1(G93A) mice

We first assessed LINGO-1 expression at different stages of the disease, at pre-symptomatic (postnatal day (P) 75), symptomatic (P110), and end-stage (P135) by Western blot analysis. However, we did not succeed in detecting any changes in expression levels (Figure 1A, B). Due to small litter sizes and the short time window for animal reproduction, it was challenging to obtain enough animals to perform a comprehensive study including both sexes. However, at the late stage (P135), we were able to compare expression levels between males and females (Figure 1C, D), and again, no differences were detected. Considering that Western blot analysis allows the determination of overall expression of LINGO1 in the spinal cord samples, but that this approach may be unsuitable to detect restricted expression variations in specific locations of the tissue, we then performed immunohistofluorescence experiments, enabling a more detailed analysis.

**Figure 1:**
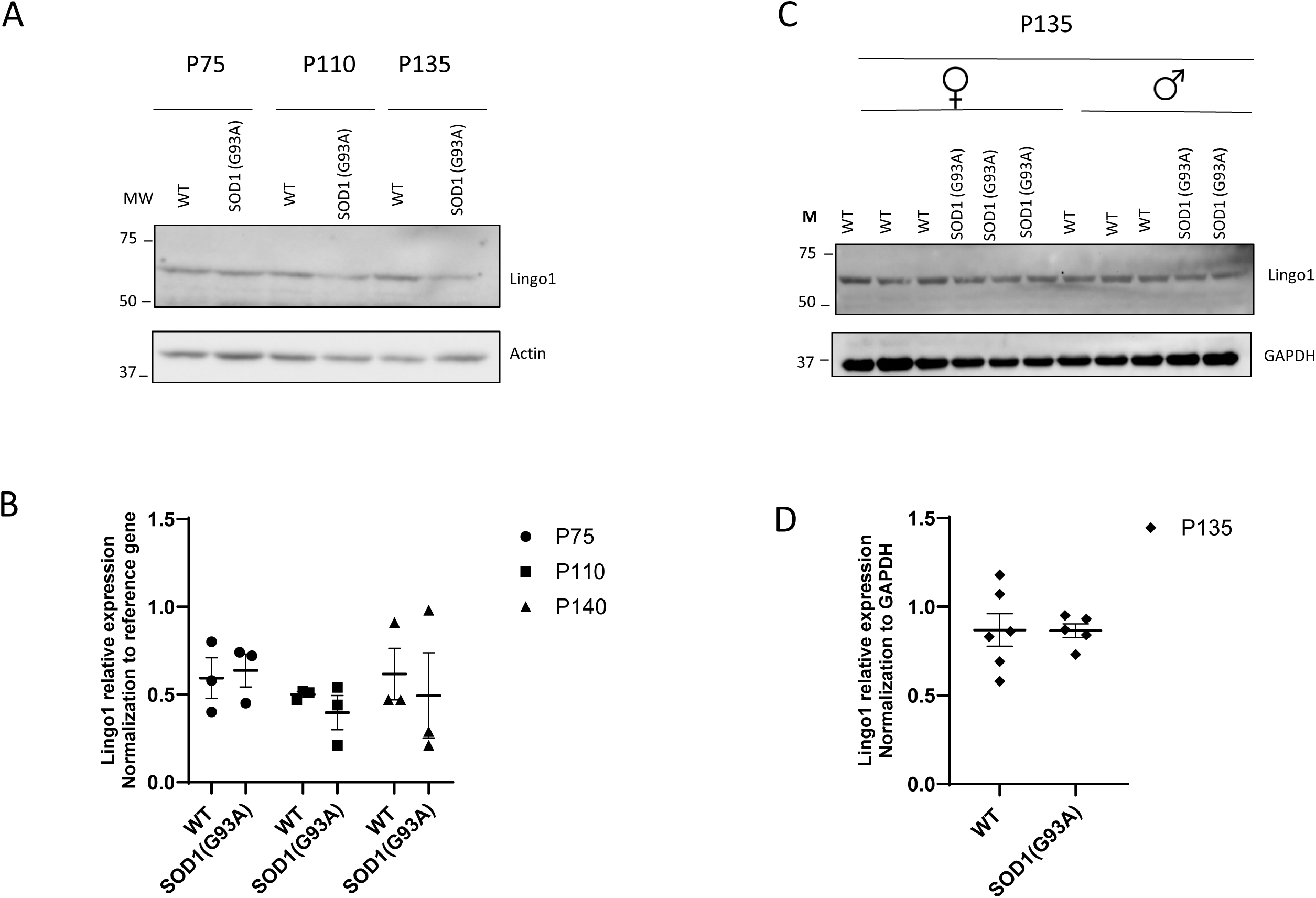
LINGO-1 expression in male and females by western blot analysis. **(A)** Representative western blot analysis of LINGO-1 expression in spinal cord slices derived from from wild-type animals (WT), pre-symptomatic (P75), symptomatic (P110) and end (P135) stages SOD1 (G93A) mice are shown and **(B)** normalized with actin expression. **(C)** Representative blot showing LINGO-1 expression in both males and females at end **(P135)** stages SOD (G93A) mice are shown and compared with labelling from wild-type animals (WT) and **(D)** normalized with reference gene expression. Values are means ± SEM. Two-way ANOVA was used for statistical analyses followed by Sidak’s multiple comparisons test and there is no significant difference between groups.

Therefore, we analyzed LINGO1 expression in the spinal cord of the SOD1(G93A) mutant and age-matched wild-type control mice using immunolabeling on tissue slices collected at the same different stages of the disease, P75, P110 and P135. Again, due to small litter sizes, the short time window for animal reproduction and the need to keep alive transgenic males for reproduction, we decided to perform immunostaining experiments only in females. (Fig. 2A-G). In the wild-type mice, LINGO1 was faintly detectable across all time points in the white matter while a clear LINGO1 expression could be observed in large cell bodies in the grey matter of the ventral horn of the spinal cord. In the SOD1(G93A) mice, LINGO1 expression was significantly increased in the white matter at P110 and to a lesser extent at P135 compared to the wild-type animals (Fig. 2A-D). Similarly, a higher LINGO1^+^ immunofluorescence could be observed in the grey matter at both P110 and P135 SOD1 (G93A) mice (Fig. 2A-C, F). In the white matter, the filamentous shape of the LINGO1^+^ labeling suggested that LINGO1-expressing cells could be astrocytes. In agreement with this hypothesis, co-labeling of the slices with the astrocyte marker GFAP confirmed LINGO1 and GFAP colocalization in the white matter. In addition, quantification of the overlapped LINGO1^+^GFAP^+^ signals indicated a significant increase of the double immunofluorescent signal in P110- and, at much lower extent, P135 SOD1 (G93A) (Fig. 2A-C, H, I). In the grey matter, GFAP-expressing astrocytes, which displayed a stellate shape as expected, did not co-express LINGO1 (Fig. A-C). The GFAP^+^ immunofluorescent area was nevertheless strongly increased in the grey matter in P110 and P135 SOD1 (G93A) mice (Fig. 2G) whereas it remained unmodified in the white matter of the mutant animals (Fig. 2E). The shape and size of LINGO1-expressing cells detected in the grey matter suggested that these cells were neurons. By performing a double labeling using LINGO1 and the motor neuron marker, choline acetyltransferase, we confirmed that choline acetyltransferase^+^ cells co-expressed LINGO1 (Fig. 2J). However, additional LINGO1^+^ cells, which displayed a similar morphology, did not co-express choline acetyltransferase, suggesting they could correspond to other neuronal subpopulations (Fig. 2J). The total number of LINGO1^+^ cells as above found for the LINGO1^+^ fluorescent area), was increased in the grey matter from the P110 and P135 mutant mice compared to age-matched wild-type animals (Fig. 2A-C, F, J, K). Thus, LINGO1 expression is significantly increased both in the grey and white matter of the spinal cord of symptomatic and end-stage SOD1 (G93A) mice, mostly in astrocytes in the white matter and in neurons in the grey matter, two cell types known to be involved in the regulation of the myelin regeneration process.

**Figure 2.**
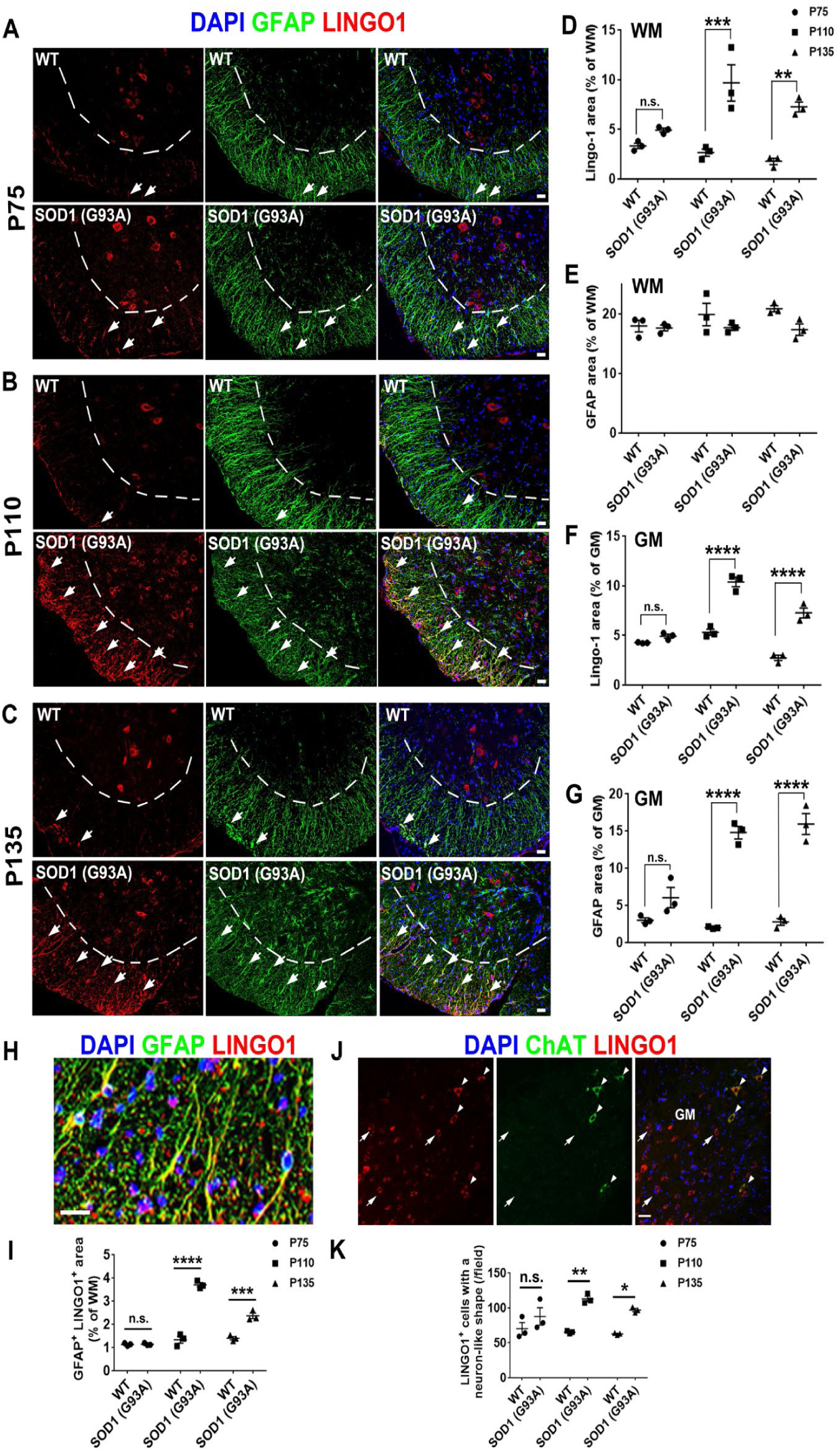
LINGO1 expression is increased in the spinal cord from SOD1 (G93A) mice at symptomatic and end stages of the disease. **(A-C)** Representative LINGO1 (red) and GFAP (green) immunohistofluorescent labelling in spinal cord slices derived from pre-symptomatic (P75), symptomatic (P110) and end (P135) stages SOD1 (G93A) mice are shown and compared with the labelling detected in the spinal cord from wild-type (WT) animals. The dotted lines delineate the border between the gray (top) and the white (bottom) matter in the ventral spinal cord. **(D-G)** Quantification of the LINGO1 and GFAP immunofluorescent signals derived from n=3 female mice per genotype and per age in the gray matter (GM) and in the white matter (WM). **(H, I)** High magnification of LINGO1 (red) and GFAP (green) double immunostaining in the white matter of P135 SOD1 (G93A) spinal cord (H) and quantification of the overlapped labelling (yellow) in P75, P110 and P135 mice (I). **(J, K)** Double LINGO1 (red) / choline acetyltransferase (ChAT, green) immunostaining in the grey matter of P110 SOD1 (G93A) spinal cord. The white arrowheads show LINGO1^+^ ChAT^+^ motor neurons while the white arrows indicate LINGO1^+^ ChAT^-^ cells with a neuron-like shape. Values are means ± SEM. Two-way ANOVA was used for statistical analyses followed by Sidak’s multiple comparisons test. *, *p* < 0.05; **, *p* < 0.01; ***, *p* < 0.001; ****; *p* < 0.0001 compared to age-matched WT animals; n.s., non-significant. Scale bars: 50 µm.

### miR-138 is dysregulated at pre-symptomatic stage of the disease

Given the increased protein-level expression of LINGO1 in the spinal cord of symptomatic and end stage SOD1 mutant mice, we investigated whether this increase could be correlated with upregulated mRNA levels. RT-qPCR analysis of LINGO1 transcripts did not show any differences between genotypes (Figure 3A), suggesting that the observed protein-level changes might be post-transcriptionally regulated.

**Figure 3.**
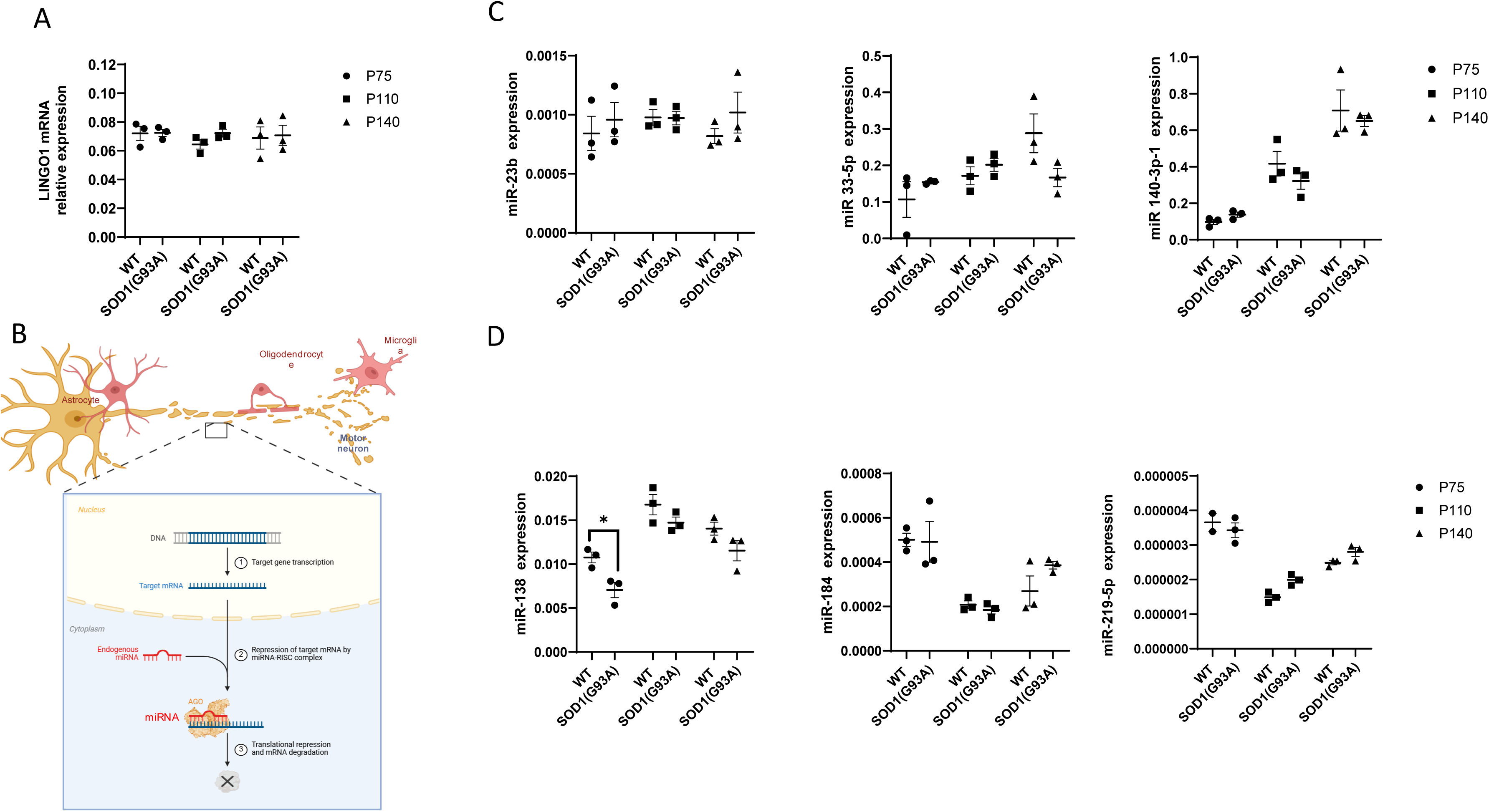
LINGO1 and miRNA expression in spinal cord of SOD1(G93A) at different stages of the disease compared to control mice (WT). **(A)** Expression of *Lingo1* in spinal cord in qPCR from WT and SOD1 (G93A) mice at P75, P110, P140 **(B)** Schematic overview of microRNA functions in neurodegenerative that silences gene expression by translational inhibition or target RNA degradation. **(C, D)** Representative graphs of RT-qPCR analysis of miRNA in spinal cord from *n* = 3 animals (both male and female). Reference genes used are: *Ppia*, *Hprt1* and *Gapdh*. Values are means ± SEM from *n* = 3-5 animals. Two way ANOVA was used for statistical analyses followed by Sidak’s multiple comparisons test. *, *p* < 0.05.

To explore this hypothesis, we examined a classical mechanism of post-transcriptional gene regulation, the miRNA-mediated inhibition of translation (Figure 3B). We focused on three miRNAs predicted to regulate LINGO1 transcription/translation, miR-23b, miR-33-5p, and miR-140-3p. We broadened our analysis to include additional miRNAs involved in myelination, miR-138, miR-184, and miR-219-5p. We quantified the expression levels of these miRNAs by RT-qPCR in control and SOD1 (G93A) mice across three disease stages. We did not observe any significant differences between genotypes for the LINGO1-targeting miRNAs (Figure 3C). However, we detected a significant decrease in miR-138 expression in SOD1 (G93A) mice at early disease stage (Figure 3D).

### Early postnatal oligodendrogenesis is impaired in the SOD1 (G93A) mutant

Since miR-138 inhibits factors that suppress OL differentiation [40], we then focused on the unexplored effects of SOD1 (G93A) mutation on perinatal oligodendrogenesis. For this purpose, we analyzed OPC production in the dorsal forebrain, thus distal from the motor neurons that further degenerate in the adult mutant. We focused on the ventricular-subventricular zone (V-SVZ), an extensive germinal layer lining the lateral ventricles, that is critical for oligodendrogenesis and post-natal forebrain myelination [33]. We traced newly generated OPCs originating from the V-SVZ at birth (P0) and at P5, a peak period of OPC production. At P0, the total number of PDGFRα^+^ OPCs did not differ between genotypes (Fig. 4A, B), but their capacity to proliferate was significantly decreased in mutant mice (Fig. 4A, D). At P5, SOD1(G93A) mice displayed a significant reduction in total OPC number (Fig. 4B), proliferating OPCs (Fig. 4C), and OPC proliferative capacity (Fig. 4D). These findings indicate impaired OPC generation as soon as the perinatal period in SOD1 (G93A) mutant mice.

**Figure 4.**
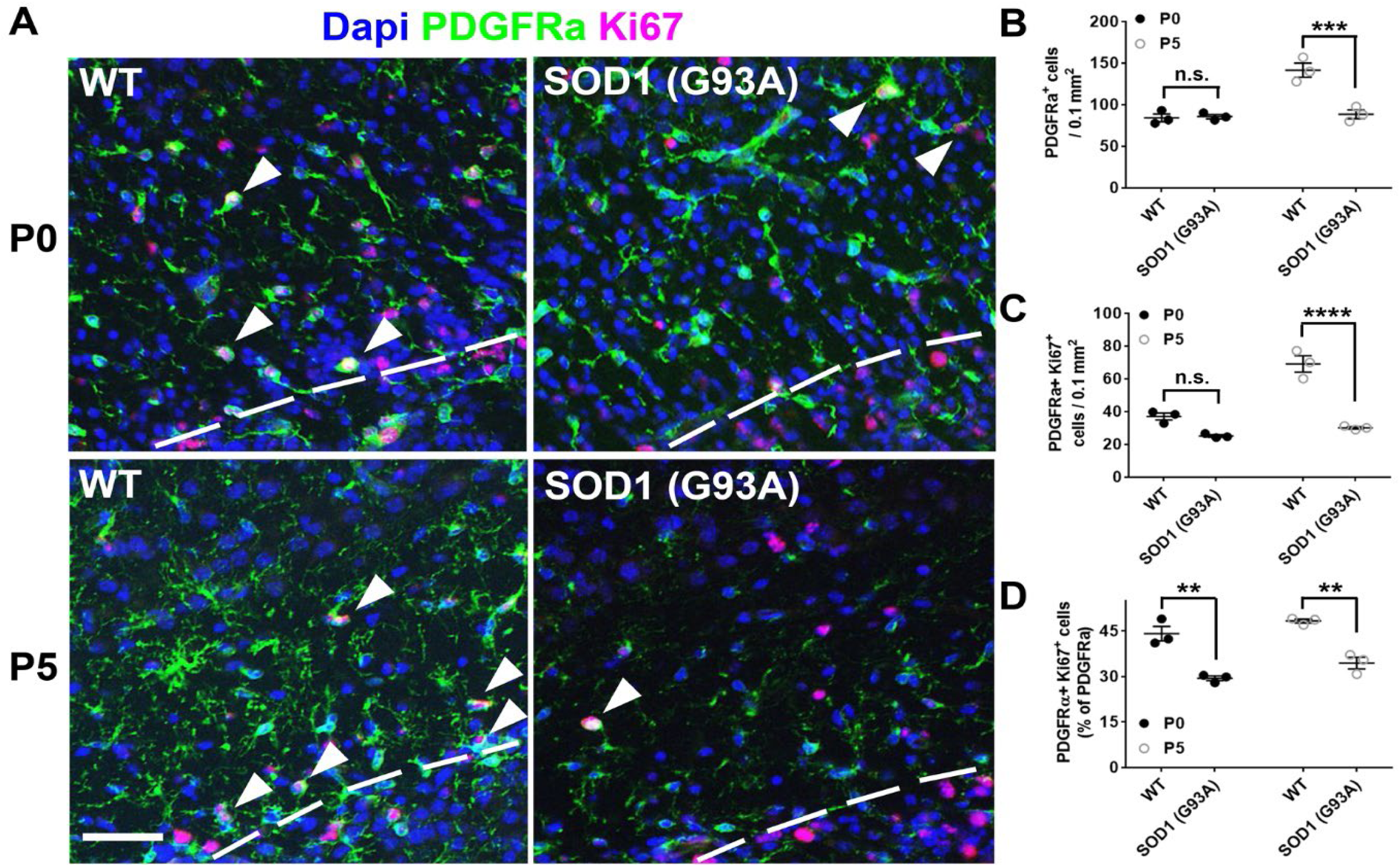
Oligodendroglial cells are generated at a much lower level in the SOD1 (G93A) mutant than in the WT animals at the perinatal period. **(A)** Slices derived from WT or SOD1 (G93A) mutant mice, observed at the level of the forebrain at the indicated postnatal age (P0, P5) and immunostained by using PDGFRα and Ki67 antibodies. The dotted line delineates the border between the germinative zone (bottom) and the developing corpus callosum (top). The white arrowheads indicate double-labelled cells. (**B-D)** Quantification of the density of PDGFRα^+^ OPCs (B), PDGFRα^+^ Ki67^+^ proliferating OPCs (C) and the proportion of OPCs able to proliferate (D). Two-way ANOVA was used for statistical analyses followed by Sidak’s multiple comparisons test. **, *p* < 0.01; ***, *p* < 0.001; ****, *p* < 0.0001; n.s., non-significant. Scale bars: 50 µm.

### miR-138 transcription is dysregulated at the perinatal period in SOD1 (G93A) mutant in the opposite way compared to adulthood

Given the established roles of LINGO1 and miR-138 in oligodendrocyte proliferation and differentiation during development, we next evaluated their expression at early postnatal stages. However, as already observed, when we evaluated the expression of LINGO1 by western blot analysis, we did not observe any changes in LINGO1 transcription in whole brain of SOD1 (G93A) compared to wild-type animals (Figure 5A, B). In contrast, we show that miR-138 transcription is increased at P8 compared to wild-type controls (Figure 5C), which may suggest higher repression of the inhibitors of oligodendrocyte differentiation and higher promotion of oligodendrocyte maturation. Thus, in response to the decreased number of OPCs (Figure 4B), miR-138 up-regulation suggests a compensatory mechanism to inhibit molecular repressors of myelination and potentially promote OPC maturation.

**Figure 5.**
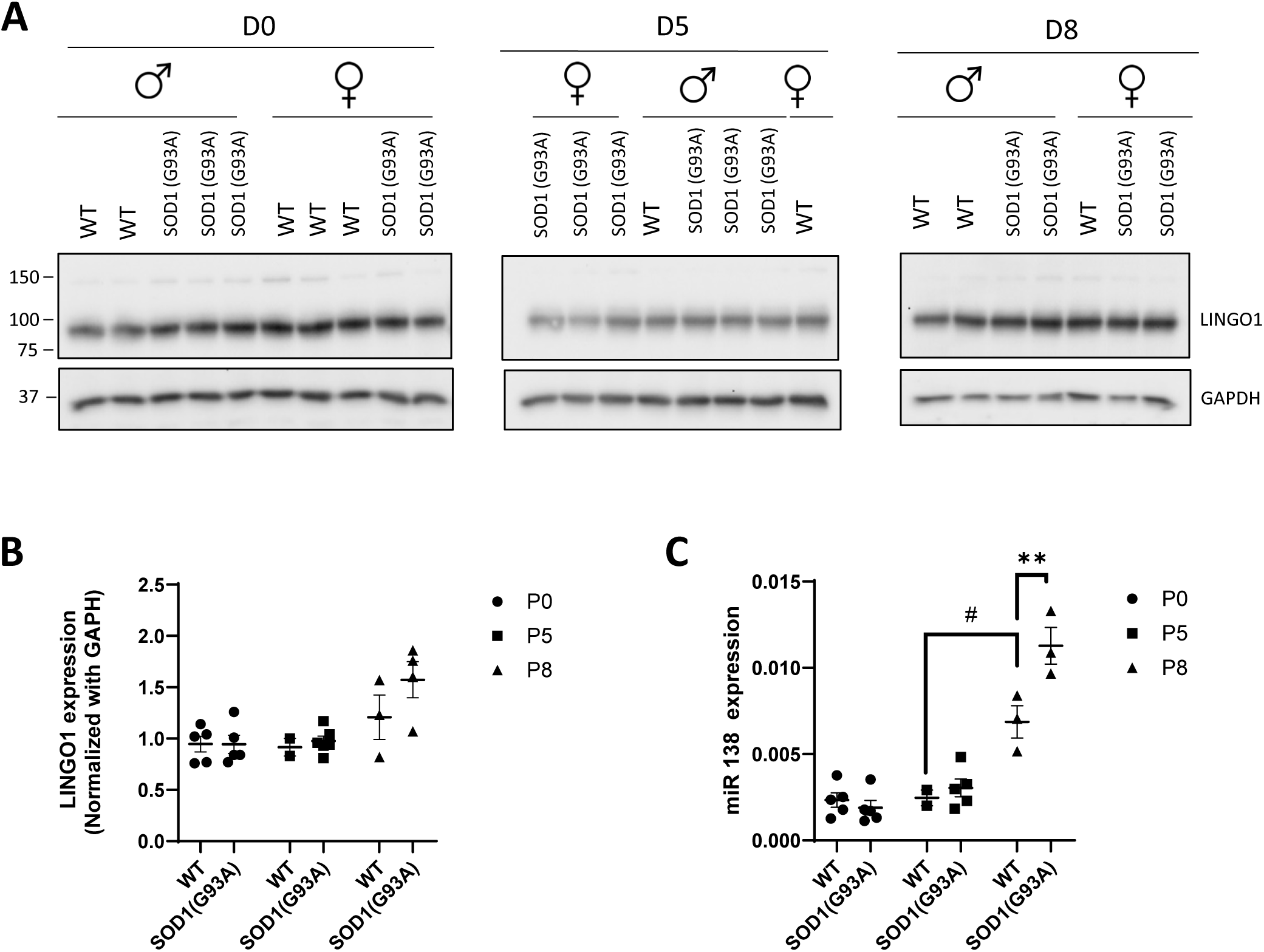
Expression of LINGO1 and miR-138 in brain from wild type WT and SOD1 (G93A) mice during the early postnatal development. (A) Expression of LINGO-1 was performed by western blot analysis in brain of WT and SOD1 (G93A) mouse at P0, P5, P8 days and (B) was normalized with GAPDH expression, (C) Representative graph of RT-qPCR analysis of miRNA in brain of WT and SOD1 (G93A) mouse at P0, P5, P8 days. Reference genes used are: *Ppia*, *Hprt1* and *Gapdh*. Values are means ± SEM from *n* =3-5 animals. Two-way ANOVA was used for statistical analyses followed by Sidak’s multiple comparisons test. **, *p* < 0.01; versus WT at the same stage; #*p* <0.05 versus WT at P5.

## Discussion

LINGO1 is a negative regulator of OPC differentiation and myelination, and its dysregulation is frequently associated with impaired remyelination. During development, LINGO1 expression has been reported in neurons and oligodendroglial cells, but not in astrocytes [14]. LINGO1 expression was also found to peak during early OPC specification, and then to decline allowing OPC differentiation and maturation into myelinating OL [34]. Consistently, preventing LINGO1 function has been successful in promoting recovery in inflammatory and non-inflammatory animal models of demyelination [35, 36].

Interestingly, our findings in the SOD1(G93A) ALS mouse model reveal an increase in the expression of LINGO1 in adulthood, at symptomatic disease stages, namely in subpopulations of astrocytes and neurons. Previous works have reported that the endogenous expression of LINGO1 in neurons [37] or its ectopic expression in cultured astrocytes [38] are both able to inhibit oligodendrocyte differentiation. Therefore, the increased expression of LINGO1 that we observed in the SOD1 (G93A) mutant at the late-stage of the disease might contribute to the failure of myelin maintenance and repair during neurodegeneration, besides the concomitant progressive degeneration of oligodendrocytes previously reported in the SOD1 (G93A) model [4].

However, the upregulation of LINGO1 at protein levels is not associated with changes in LINGO1 transcription. Although mRNA levels are often used as a proxy for gene expression, numerous studies have demonstrated that transcript abundance modestly correlates with protein levels. This uncoupling is well documented across tissues, including the nervous system, and indicates that protein levels can be regulated by mechanisms independent of transcriptional control [39]. In our study, unchanged levels of LINGO1 mRNA, combined with unchanged levels of predicted miRNAs targeting LINGO1, suggests that the increased protein expression in SOD1 mutant mice does not result from transcriptional or canonical miRNA-mediated post-transcriptional regulation. Instead, alternative mechanisms may account for this discrepancy. First, translation efficiency can vary independently of mRNA abundance. For instance, altered ribosomal loading and translational initiation factors [40] may selectively enhance LINGO1 translation. Second, protein stability is frequently altered in pathological contexts. Indeed, reduced proteasomal or lysosomal degradation, or disease-induced post-translational modifications (*e.g*., phosphorylation or ubiquitination shielding), can markedly increase protein levels without requiring increased mRNA synthesis [41]. Third, ALS is associated with widespread proteostasis disturbances, leading to selective accumulation of certain membrane or signaling proteins despite unchanged transcript levels [42]. Thus, the increased LINGO1 protein expression observed at ALS symptomatic stages, likely reflects dysregulation at the translational or post-translational level, rather than changes in gene transcription or miRNA-mediated repression. This raises the possibility that increased expression of LINGO1 may promote its multimerization, given its well-documented propensity to form dimers and higher-order assemblies [38, 43–45]. Such a multimeric organization could amplify or prolong LINGO1–associated inhibitory signaling, particularly during oligodendrocyte differentiation. Nevertheless, the oligomeric state of LINGO1 under these conditions warrants further investigated.

Moreover, our data reveal a significant increase in miR-138 expression during early postnatal development (from P5 to P8), which coincides with the critical window of OPC differentiation and initiation of myelin protein synthesis. MiR-138 is a pro-myelinating microRNA that promotes OPC differentiation by repressing transcriptional inhibitors [46, 47]. In the healthy developing CNS, miR-138 expression rises during the transition from OPCs to pre-myelinating OLs and remains elevated as cells acquire myelinating identity making this microRNA a facilitator of myelin gene induction. Interestingly, in the SOD1 mutant mice, we observed an increase in miR-138 expression at P8 that could reflect an endogenous mechanism of compensation aimed at counteracting the impaired oligodendrogenesis that we observed in this mutant. This observation contrasts with the data obtained at presymptomatic stages of the disease, namely a significant reduction in miR-138 expression, which suggests that disease progression disrupts the regulatory balance required for effective oligodendrocyte maturation. The reduced expression of miR-138 at P75 might alleviate the inhibitory effects of this miRNA on anti-myelination pathways thus driving myelin impairment, in a manner that contrasts with the early postnatal conditions. Interestingly, because this downregulation occurs prior to appearance of clinical symptoms, miR-138 may represent a presymptomatic biomarker candidate of oligodendroglial vulnerability in ALS, a hypothesis that would deserve to be further investigated.

To conclude, our findings highlight the increase in the number of fibrous astrocytes and neurons that express LINGO1 in the spinal cord of SOD1 (G93A) mice, which might be consistent with the contribution of LINGO1 to the demyelination characterizing the disease. Moreover, we show a dual and complementary regulation of LINGO1 and miR-138 across disease progression, likely pointing toward a complex but interpretable balance between inhibitory and pro-myelinating forces in ALS. During early postnatal development of the disease, we reveal an impairment of oligodendrogenesis, which is temporally associated with an increase in miR-138, a pattern consistent with a possible compensatory mechanism by which miR-138 upregulation might promote the repression of anti-myelin transcriptional programs and subsequently OPC differentiation and myelination. However, as disease progresses toward presymptomatic stages, this balance shifts. miR-138 expression falls significantly, possibly reducing its impact on pathways known to inhibit myelination. In parallel, we report that LINGO1 expression rises at later symptomatic stages, converging toward the inhibitory profile observed in classical models in which remyelination fails. Therefore, the SOD1 mutant mice might display a dual and temporally dependent phenotype, namely a compensatory phenotype during development and an inhibitory phenotype at the late stage of the disease, which might contribute to the progressive failure of myelin maintenance and repair in ALS.

## Acknowledgments

Some schematics were created with BioRender.com and used under an academic individual License. This work was supported by Orléans University (ATER position for Julie Bourseguin and Mireille Moussaed) and by the association pour la recherche sur la SLA, ARSLA, to E. Traiffort (Grant # RAK19097LLP).

## Author contributions: CRediT

Conceptualization : Amina Zahaf, Julie Bourseguin, Elisabeth Traiffort and Séverine Morisset-Lopez

Methodology and Investigation : Amina Zahaf, Julie Bourseguin, Laetitia Cobret, Abdelmoumen Kassoussi and Mireille Moussaed

Validation: Amina Zahaf, Julie Bourseguin, Laetitia Cobret, Abdelmoumen Kassoussi, Mireille Moussaed, Cédric Raoul, Elisabeth Traiffort and Séverine Morisset-Lopez

Writing-original draft: Séverine Morisset-Lopez and Elisabeth Traiffort

Funding acquisition and Supervision: Séverine Morisset-Lopez and Elisabeth Traiffort

## Ethics approval

All animal procedures were conducted in accordance with the European Union Directive 2010/63 and the ARRIVE guidelines. Experiments on mice were authorized by the French Ministry of Teaching, Research and Innovation (APAFIS #6224-2016072711103977v3; APAFIS#30104-2021022609497819 and APAFIS #40248-2022122909437846).

